# Probing apoptosis signaling proteins in single living cells for precision efficacy evaluation of anti-cancer drugs

**DOI:** 10.1101/663013

**Authors:** Yanrong Wen, Jia Liu, Hui He, Zhen Liu

## Abstract

*In vitro* efficacy evaluation is critical for anti-cancer drug development and precision cancer treatment. Conventional methods, which mainly rely on cell viability and large cell populations, suffer from apparent disadvantages, such as signaling pathway-nonspecific, failing to reflect the real sensitivity of the patient, tedious and time-consuming. Herein, we present a new analytical tool, termed single-cell plasmonic immunosandwich assay (scPISA), for precision efficacy evaluation of anti-cancer drugs. It allows for facilely probing individual signaling proteins as well as protein-protein complexes in single living cells. Based on this approach, two apoptosis signaling proteins, cytochrome C (Cyt C) and caspase-3, were proposed as apoptosis indicators, while three new parameters were proposed as criteria for quantitative efficacy evaluation. Using two typical cytotoxic drugs, actinomycin D (Act D) and staurosporine (STS), as model drugs, the evaluation was found to be consistent among the indicators and parameters. Metformin, a potential anti-cancer drug, was then evaluated using this approach. Interestingly, metformin alone was found to be a less effective anti-cancer drug but its combination with Act D dramatically improved the overall efficacy. The scPISA approach exhibited several unique strengths over conventional assays, including comprehensive and self-consistent evaluation, signaling pathway-specific, simple procedure and high speed. Thus, it holds great promise for personalized drug screening and precision medicine.

---

*In vitro* evaluation of drug efficacy is critically important for anti-cancer drug development and precision cancer treatment^1–3^. Conventional tools for assessing drug efficacy, such as MTT assay and clonogenic assay, rely on cell viability as well as a large cell population or cell culture^4–7^. An apparent limitation of cell viability-based approaches is that the molecular mechanism underlying the effects of the drugs is lacking. To overcome this issue, signaling pathway-based drug efficacy evaluation has been gained increasing attentions^8–12^. On the other hand, large cell population-based approaches also suffer from apparent drawbacks. These approaches fail to reflect the real sensitivity of the patient’s cancer cells, because the cell number from a patient is usually limited whereas cultured cancer cells are not representative of those in the patient^13^. Besides, these assays measure the average response of a cell population across a long period of drug stimulation and thereby fail to report any intermediate events of drug action^14^. Moreover, these approaches are tedious and time-consuming. To solve these issues, single-cell analytical approaches have been developed for the evaluation of drug efficacy^15–19^. However, single-cell analytical approaches that permit determination of signaling proteins in single cells still remains limited, especially in single living cells^20–22^. This is mainly due to the fact that many signaling pathway proteins are present at very limited abundance particularly at early stages^23,24^. Since there are no PCR-like tools for protein amplification, an ultrasensitive detection is critical for the detection of pathway proteins in early signaling events.

Herein, we present a new analytical approach, called single-cell plasmonic immunosandwich assay (scPISA), for precision efficacy evaluation of anti-cancer drugs through probing apoptosis signaling proteins in single living cells. This approach mainly relies on our previous PISA approach that permitted for facile determination of low-copy-number proteins (less than 1000 copies per cell) in single living cells with single-molecule sensitivity^25^. Since the PISA approach has also been applied to the detection of protein biomarkers in living animals^25^ and other biological samples such as urine^26^ and serum^27^, the PISA approach herein is particularly termed as scPISA. The approach mainly relies on the combination of *in-vivo* immunoaffinity extraction in single living cells and ultrasensitive plasmonic detection (**Scheme 1**). An extraction probe immobilized with an antibody against a signaling protein of interest was precisely inserted into a single apoptotic cell of interest through a micromanipulator. After extraction for a short period of time, the probe was taken out of the cell and washed to remove unwanted species, followed by labeling the captured species with Raman labeling nanotags immobilized with another antibody specific to the same signaling protein or a second signaling protein with which the former one may bind. Sandwich-like complexes formed on the extraction probe were detected by a Raman spectrograph. This approach allowed for facilely probing individual signaling proteins as well as protein-protein complexes in drug-induced apoptosis. Based on this single-cell analytical tool, we proposed a new strategy for quantitative evaluation of anti-cancer drug efficacy. Two key signaling proteins of apoptosis, cytochrome C (Cyt C) and caspase-3, were adopted as apoptosis indicators, while a set of new parameters calculated from the time- and dose-dependent response curves were proposed as criteria for quantitative evaluation of anti-cancer drug efficacy. Using two typical cytotoxic drugs, actinomycin D (Act D) and staurosporine (STS), as model drugs, the apoptosis evaluation parameters were measured and found to be consistent with each other. Using the proposed approach, the anti-cancer efficacy of metformin used alone or in combination with Act D was evaluated. The results suggested that metformin alone was not a very effective anti-cancer drug; however, its combination with Act D dramatically improved the overall efficacy. Comparison with MTT assay revealed that the scPISA approach provided comprehensive and more reliable evaluation. As compared with conventional assays, this new approach exhibited several unique strengths, including comprehensive and self-consistent evaluation, signaling pathway-specific, simple procedure and high speed. Thus, this approach opened a new access for precision evaluation of anti-cancer efficacy. It holds great promise in personalized drug screening and precision medicine.

**Scheme 1.**
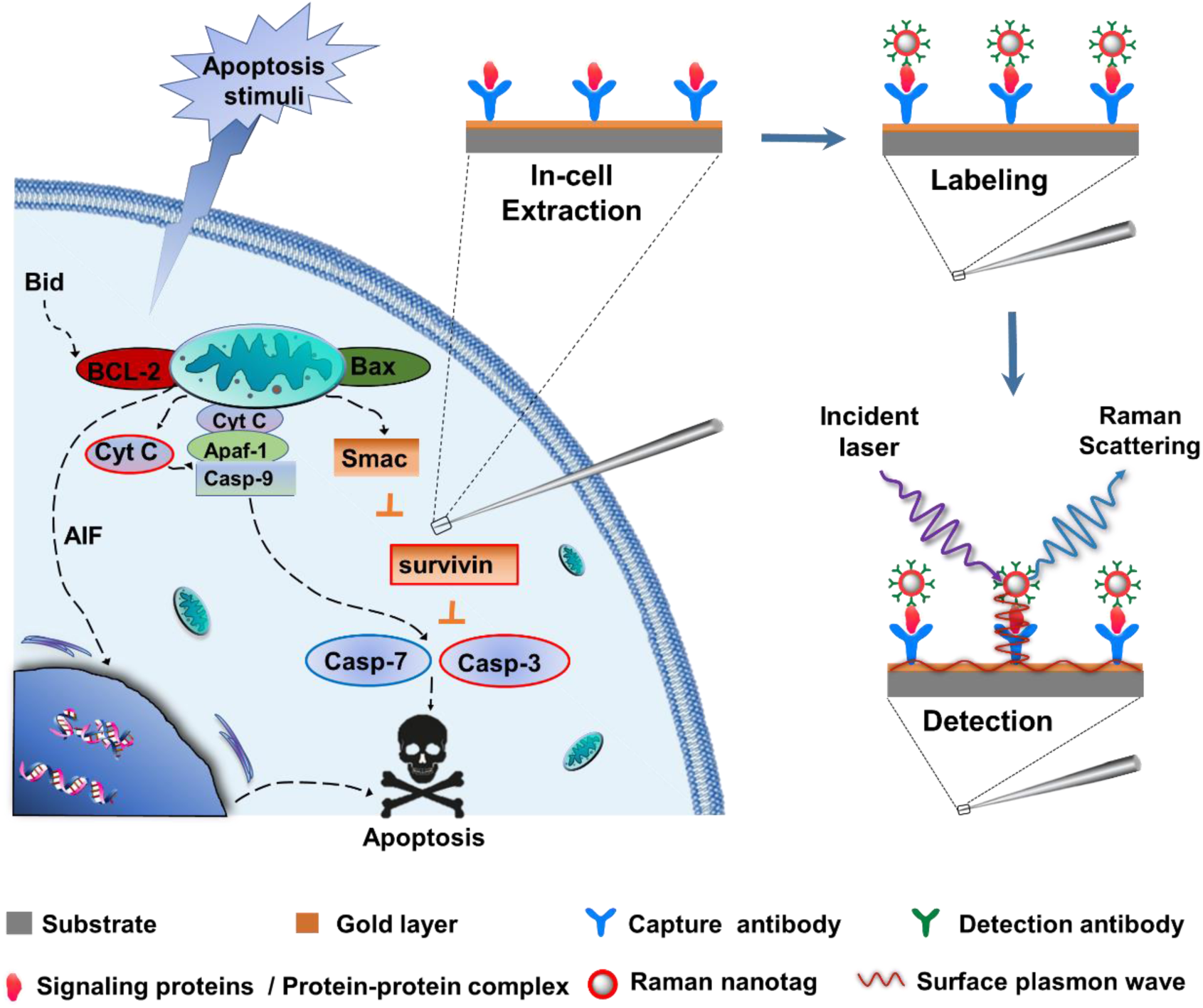
Schematic diagram of probing the signaling proteins of apoptosis in single living cells via scPISA.

## RESULTS

### Characterization and verification

Extraction microprobes and Raman nanotags for scPISA were prepared according to the procedures shown in **Supplementary Fig. 1**. The obtained extraction probes exhibited tiny tips of ca. 0.6-1 μm in diameter **(Fig. 1a)** and limited background Raman signal (**Supplementary Fig. 2a**). The antibody-immobilized Raman nanotags showed a core-size of ca. 60 × 80 nm (**Fig. 1b**) and significant Raman signal (**Supplementary Fig. 2b**). The intensity of the peak at 1,435 cm^−1^ was employed for the assays of all signaling proteins and protein-protein complexes. Dynamic light scattering (DLS) characterization further confirmed the core-size of the Raman nanotags, while UV-vis absorption spectra showed that silica encapsulation and antibody immobilization did not significantly change the particle size since the localized surface plasmon resonance (LSPR) peak only slightly red-shifted (**Supplementary Fig. 3**). By using the prepared extraction microprobes and Raman nanotags, the developed scPISA approach was applied to the detection of Cyt C and caspase-3 in solution and apoptotic single Hela cells. The Raman intensities for apoptotic cells changed from cells to cells while those for the standard solutions were quite reproducible (**Fig. 1c and Fig. 1d**). The signal variation was attributed to the microheterogeneity of cells. The dependence of the Raman intensity on the concentration of Cyt C and caspase-3 in standard solutions was investigated. As shown in **Fig. 1e and Fig. 1f**, it obeyed a typical logarithmic function; that is, the Raman intensity increased with the logarithm of the protein concentration, and the dependence was linear within a certain range of concentration. The linear range spanned 5 and 6 orders of magnitude for Cyt C and caspase-3, respectively. The limit of detection (LOD) of this approach was 7.2 ×10^−14^ M for Cyt C and 5.8 ×10^−15^ M for caspase-3 (S/N > 3). Such ultrahigh sensitivity enabled determining apoptosis signaling proteins of very limited concentration in single cells at early events of apoptosis to identify the pathway by which apoptosis has been induced.

**Figure 1.**
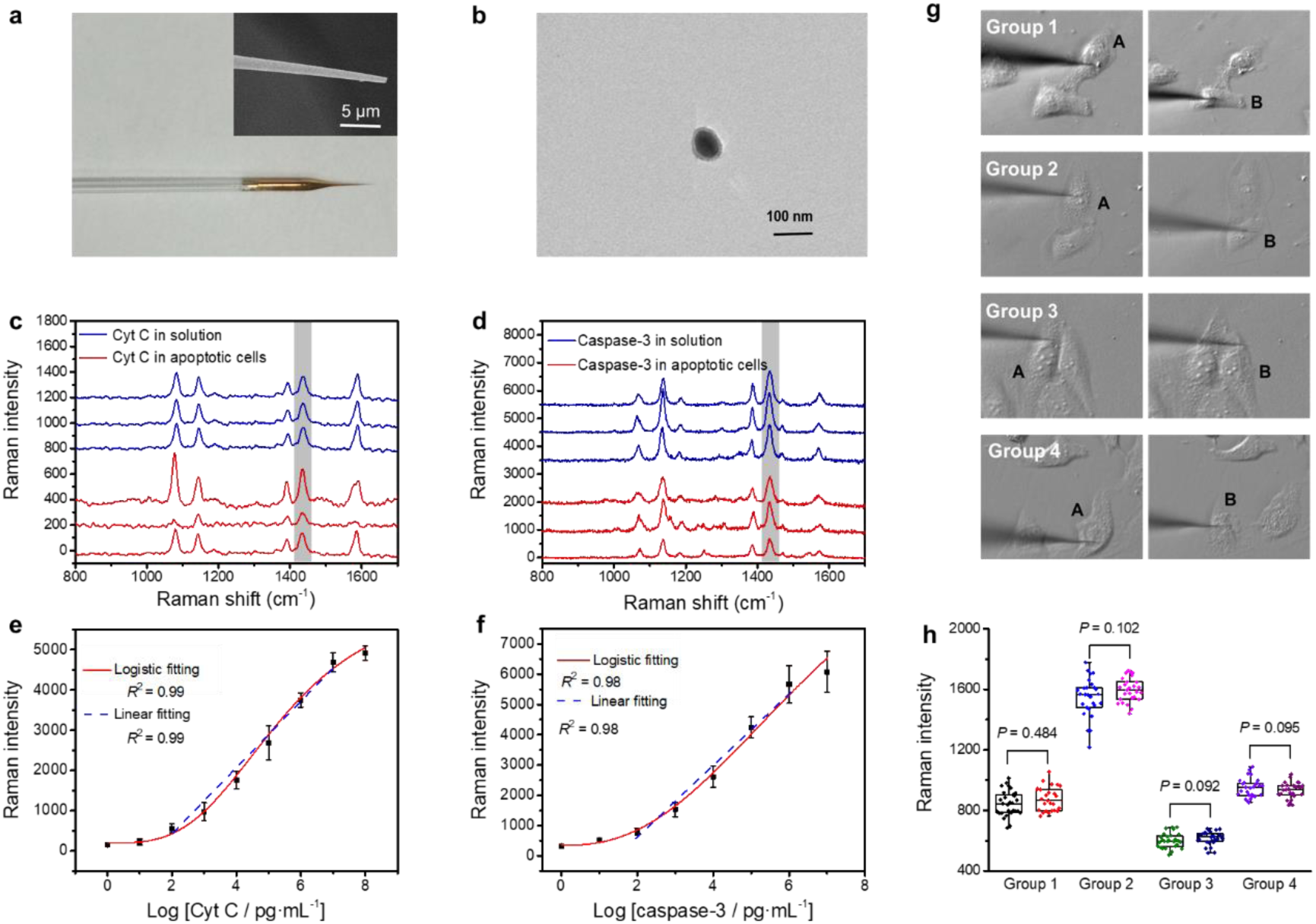
Basic characterization and verification. **(a)** Photo and SEM images of an extraction microprobe. **(b)** TEM images of Raman nanotags. **(c)** Raman spectra for Cyt C in solution (1 pg·mL^−1^) and single apoptotic Hela cells stimulated with 4 μM STS for 0.5 h. **(d)** Raman spectra for caspase-3 in solution (100 pg·mL^−1^) and single apoptotic Hela cells stimulated with 4 μM STS for 2 h. **(e)** Raman intensity-concentration dependence for **(c)** Cyt C and **(d)** caspase-3 in solution. **(g)** Images showing the insertion of extraction microprobes in 4 groups of apoptotic HepG-2 sister cells. **(h)** Box plots showing Raman intensity detected by scPISA for Cyt C in the sister cells shown in **g**.

To further examine the reliability of scPISA for probing signaling pathway proteins in single cells, Cyt C in four groups of apoptotic HepG-2 sister cells was detected. Four mother cells were first stimulated with Act D. During the induced apoptosis, each mother cell divided into two sister cells. After division, Cyt C in each sister cell was immediately probed. **Fig. 1g** shows the photos of the sister cells under investigation. Although the Raman intensity for Cyt C apparently varied among the mother cells, the intensities of the sister cells were almost the same (**Fig. 1h**). Since newly divided sister cells exhibited high similarity in their timing of apoptosis^28^, this result suggests that the scPISA approach provided a reliable tool for assessing the expression level of signaling proteins in single cells of apoptosis.

### Indicators and parameters for characterization of apoptosis

In a signaling pathway, there are a series of protein-related molecular events, in which proteins are produced, released or transformed. Clearly, it is unnecessary to monitor all signaling proteins. Rather, it is essential to monitor some key signaling proteins that can feature a specific signaling pathway. In intrinsic or mitochondrial apoptosis signaling pathway, two events are characteristic. One is the mitochondrial release of apoptogenic factors, particularly Cyt C, which is an upstream event.^29,30^ The other is the downstream caspases activation^31–33^. In this study, we used Cyt C and caspase-3 as two apoptosis indicators. Assessing the functional integrity of apoptosis signaling by simultaneous measurement of the two signaling proteins were expected to provide necessary and essential information to identify the pathway. **Fig. 2a** and **Fig. 2b** show the exposure time-dependent Raman response for Cyt C and caspase-3 in STS-induced single apoptotic Hela cells. Representative Raman spectra and the signal intensity for individual cells are shown in **Supplementary Fig. 4**. Within initial 3-4 h of exposure, the expression level of Cyt C and caspase-3 increased as increasing the exposure time. After that, the expression level of Cyt C was saturated while that of caspase-3 gradually reduced. The different time-dependent response curves were probably due to the different functions of Cyt C and caspase-3. The main function of Cyt C is to bind with associated species including apoptotic protease activating factor-1 (Apaf-1) and pro-caspase to form a protein machinery known as apoptosome, which cleaves the pro-caspase to its active form caspase-9 and finally activates the effector caspase-3^34^. While the main function of caspase-3 is to proteolytically degrade the intracellular contents. We attributed the reduction in the expression level of caspase-3 after its maximum to that the consumption speed of caspase-3 may be increased at later phase of apoptosis. In this sense, Cyt C may function as a more sensitive and stable indicator to reflect the time course of apoptosis. Fitting of the time-dependent response curve for Cyt C with the logistic function gave two parameters: 1) *I*_max_, which is defined as the maximal response of apoptosis, and 2) *T*_50_, which is the time for half maximal response. These two parameters reflect the strength and speed of apoptosis of a cell exposed to a cytotoxic drug, respectively. The dose-dependent response curves for STS-induced apoptosis are shown in **Fig. 2c** and **Fig. 2d**. Fitting of the dose-response curves with the logistic function yield a new parameter: *EC*_50_, which is defined as the concentration for half maximal response. The *EC*_50_ values for the same cell line stimulated by the same drug in terms of the expression level of Cyt C and caspase-3 separately were very close, being 1.26 and 1.22 µM, respectively. This indicates that either Cyt C or caspase-3 can function as a reasonable indicator for the dose-dependent response of single cells towards cytotoxic drug-induced apoptosis. We examined the correlation between the response for Cyt C and that for caspase-3 in the same apoptotic cells and found that there was a good correlation (**Fig. 2e** and **Fig. 2f**). This further indicates that within the same signaling pathway, the upstream and downstream signaling proteins provided equal characterization of apoptosis.

**Figure 2.**
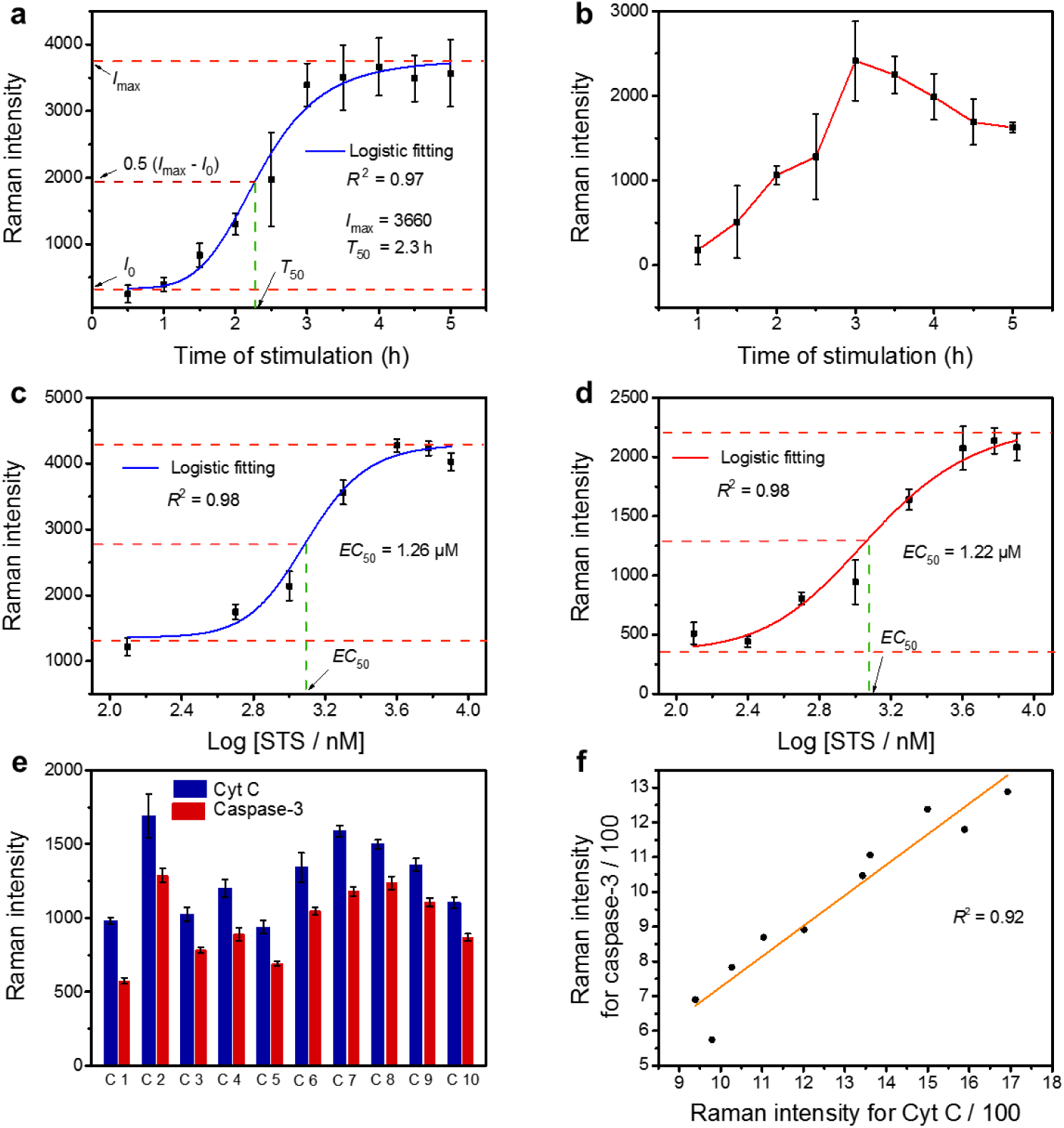
Evaluation of anti-cancer efficacy via apoptosis-related parameters. **(a)** Dependence of the average Raman intensity for Cyt C in single apoptotic Hela cells stimulated with 4 μM STS on the stimulation time. **(b)** Dependence of the average Raman intensity for caspase-3 in single apoptotic Hela cells stimulated with 4 μM STS on the stimulation time. **(c)** Dose-effect curve for STS with Cyt C as apoptosis indicator. **(d)** Dose-effect curve for STS with caspase-3 as apoptosis indicator. **(e)** Raman intensity for Cyt C and caspase-3 measured simultaneously in 10 same individual HepG-2 cells stimulated by Act D (4 μM) for 7 h. **(f)** Correlation of the Raman intensities for signaling molecule Cyt C and caspase-3 in 10 individual HepG-2 cells.

Apoptosis induced by another cytotoxic drug, Act D, using caspase-3 as the apoptosis indicator was also investigated (**Supplementary Fig. 5**). The maximal intensity was obtained at 4 h (compared to STS at 3 h) and a drop of signal response after the maximum was also observed. The three apoptosis parameters were also measured, which are listed in **Supplementary Table 1**. From the values of these parameters, we can conclude that STS is more effective than Act D in their *in-vitro* anti-cancer efficacy.

### Distinct responses of cancer and normal cells to apoptosis-inducing stimuli

We investigated the response of cancer cells and normal cells towards the stimulation of the cytotoxic drugs STS and Act D. Human normal hepatocytes cell line L-02 and hepatoma cell line HepG-2 were adopted as the model cell lines, and Cyt C was used as the indicator of apoptosis. **Fig. 3** and **Supplementary Fig. 6** show the time-dependent response in apoptotic HepG-2 and L-02 cells simulated with STS and Act D of identical concentration. **Supplementary Table 2** lists the measured *I*_max_ and *T*_50_ values. The cancer cells exhibited much higher *T*_50_ value (2.9 or 3.5-fold higher) as compared with the normal cells, though the *I*_max_ value for the former was just slightly lower than that for the latter (being 79% or 98% of the value of the latter). In other words, although nearly equal amount of Cyt C was released from the mitochondrial, the cancer cells were apparently resistant to both apoptosis-inducing drugs. Such results are in good agreement with the well-known fact that resistance to apoptosis is a hallmark of cancer cells^35,36^.

**Figure 3.**
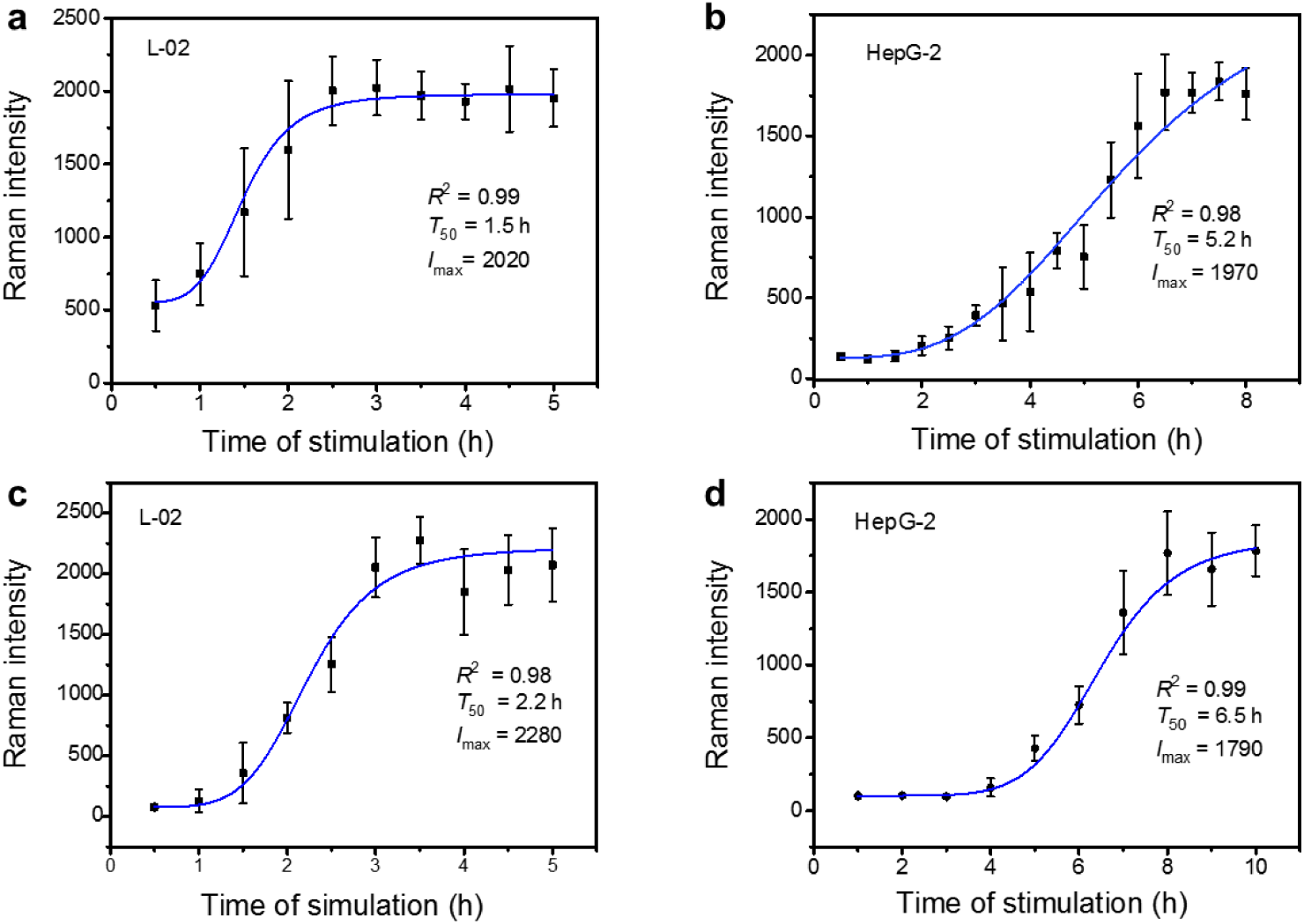
Different response of normal cells and cancer cells to anti-cancer drugs. Time-dependent Raman response for Cyt C in single **(a)** L-02 cells and **(b)** HepG-2 cells stimulated with 4 μM STS. Time-dependent Raman response for Cyt C in single **(c)** L-02 cells and **(d)** HepG-2 cells stimulated with 4 μM Act D. Error bars represent standard deviations for the average Raman intensity on 3 microprobes. The data were fitted according to logistic function.

### Probing signaling protein-protein complexes of apoptosis

Since the resistance of cancer cells to apoptosis was observed, we further investigated whether the scPISA approach can provide a reasonable explanation in molecular mechanism. Survivin, as a well-documented member of the inhibitor of apoptosis protein (IAP) family, plays a pivotal role for tumor cell survival^37,38^. Survivin inhibits caspase activation, leading to negative regulation of apoptosis^39^. It has been widely reported that survivin in cancer cells is overexpressed^25,40^. We firstly examined whether survivin can form complexes with the two signaling proteins in buffer solution. As shown in **Fig. 4a** and **Fig. 4b**, the scPISA assay revealed that survivin formed complexes with either caspase-3 or Cyt C in PBS buffer. We then investigated whether these complexes could be detected during apoptotic stimulation with STS. **Fig. 4c** and **Fig. 4d** show the time-dependent response for caspase-3-survivin complex and Cyt C-survivin complex, respectively. Clearly, despite that the signal intensity was limited, the scPISA approach permitted the detection of these complexes within apoptotic cells. Caspase-3-survivin complex was apparently detected in all apoptotic cells after the stimulation exceeded 2.5 h. As a comparison, Cyt C-survivin complex was only detected in only partial apoptotic cells within initial 2 h of stimulation. Meanwhile, it was observed that the expression level of free survivin gradually decreased as the stimulation time increased (**Supplementary Fig. 7**), which mirrors that free survivin was depleted gradually due to the formation of caspase-3-survivin complex. Thus, the reason for the resistance of cancer cells to apoptosis can be mainly attributed to the binding of survivin with caspase-3, which inhibited the activity of the latter.

**Figure 4.**
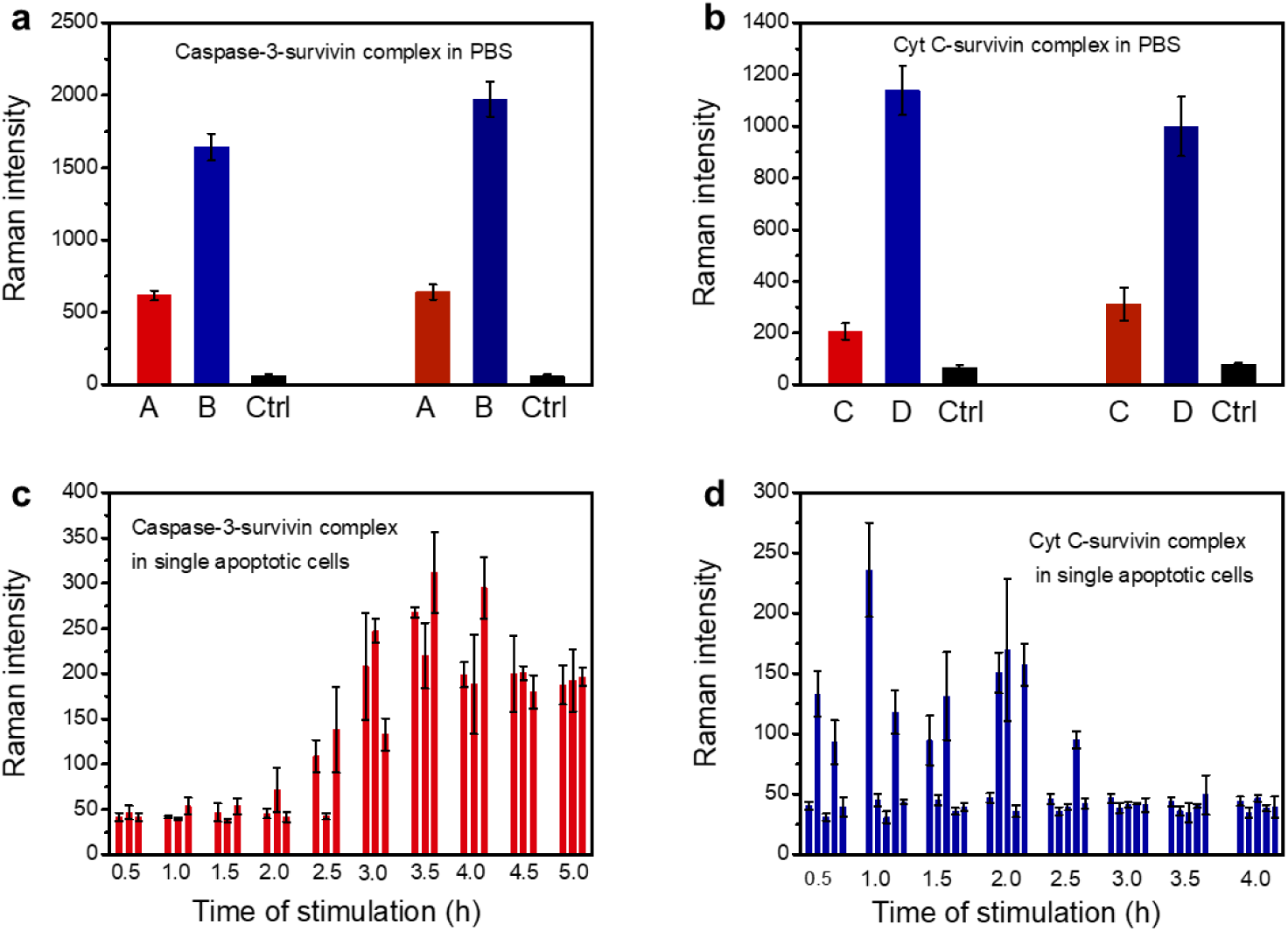
Detection of protein-protein complexes *in vitro* and in single apoptotic cells. Raman intensity for different forms of **(a)** caspase-3 and **(b)** Cyt C with survivin in PBS detected by scPISA with different extraction probe-Raman tag configurations. Raman intensity for **(c)** caspase-3-survivin complex and **(d)** Cyt C-survivin complex in single apoptotic HepG-2 cells stimulated with 4 μM STS. Error bars represent standard deviations for the average Raman intensity on 3 microprobes. A, caspase-3-survivin complex, detected by the combination of microprobes for caspase-3 recognition and Raman nanotags for survivin recognition; B, free caspase-3; C, Cyt C-survivin complex, detected by the combination of microprobes for Cyt C recognition and Raman nanotags for survivin recognition; D, free Cyt C, Ctrl used microprobes for caspase-3 / Cyt Cto extract in PBS, and labeled by Raman nanotags for survivin. The left three columns were obtained with anti-caspase-3/anti-Cyt C monoclonal antibody immobilized microprobes; the right three columns were obtained with anti-caspase-3/anti-Cyt C polyclonal antibody immobilized microprobes in **a** and **b**.

### Evaluation of apoptosis induced by metformin

Metformin, as a potential anti-cancer drug, has gained wide attention recently^41^. Although the anti-cancer mechanisms of metformin have been studied^42,43^, quantitative evaluation of its efficacy is still lacking. We applied the scPISA technology to the evaluation of apoptosis induced by metformin, through tracking the expression level of the signaling proteins in single cells. Using Cyt C as an apoptosis indicator, we firstly investigated the time course and dose-response curve for the apoptosis of single HepG-2 cells stimulated with metformin. **Fig. 5a** and **Fig. 5b** show that metformin could induce apoptosis. However, its anti-cancer effect was much weaker as compared with STS and Act D, giving a much lower *I*_max_ value and much higher *T*_50_ and *EC*_50_ values. Considering the high *EC*_50_ value, metformin might not be a practical anti-cancer drug. We then investigated the apoptosis of HepG-2 cells stimulated with the combination of metformin (10 mM) and Act D. Interestingly, through the combined use of metformin with Act D, the overall anti-cancer efficacy was dramatically enhanced, showing much higher *I*_max_ value and much lower *T*_50_ and *EC*_50_ values (**Fig. 5c** and **Fig. 5d**). **Fig. 5** shows two sets of *I*_max_ values: one is based on the time course as described above, while the other is based on the dose-response curve. Clearly, the two sets of *I*_max_ values were almost the same for identical cell line and stimulation conditions. The apoptosis evaluation parameters for metformin alone and along with Act D are listed in **Supplementary Table 3**. Meanwhile, the *IC*_50_ values measured by MTT are also listed for comparison. Overall, the *IC*_50_ values were apparently higher than the *EC*_50_ values. This was attributed to the poor sensitivity of MTT assay as well the use of much larger population of cells.

**Figure 5.**
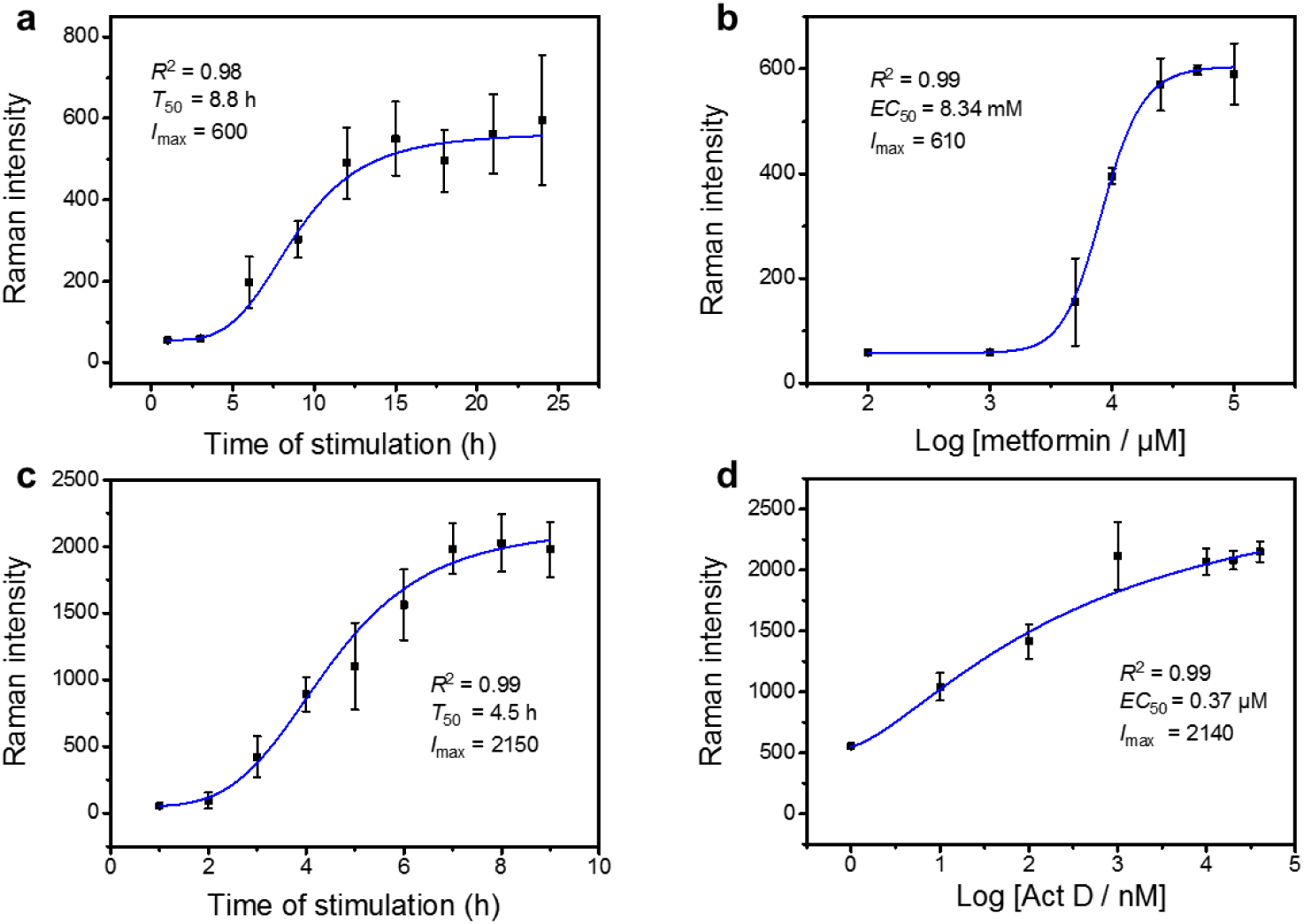
Evaluation of apoptosis inducement efficacy of metformin used alone (a and b) and in combination with Act D (c and d). **(a)** Time-dependent Raman response for single apoptotic HepG-2 cells stimulated with 10 mM metformin, using Cyt C as an apoptosis indicator. **(b)** Dose-dependent Raman response for Cyt C in single apoptotic HepG-2 cells. The data were fitted according to logistic function. **(c)** Time-dependent Raman response for Cyt C in single HepG-2 cells stimulated with combination of 4 μM Act D and metformin (10 mM). **(d)** Dose-effect curve for apoptosis induced by the combination of Act D and metformin (10 mM) using Cyt C as an apoptosis indicator.

## DISCUSSION

In this study, we first demonstrated the powerfulness of scPISA for probing apoptosis signaling proteins in single living cells. The scPISA approach exhibited several significant advantages over conventional methods for signaling pathway study, particularly Western blotting. First and the most importantly, it permitted facile single-cell dissection of not only individual cellular proteins but also protein-protein complexes in apoptosis signaling pathway. Second, the scPISA approach is straightforward, taking only a few steps. Since this approach has certain spatial resolution, it allowed for extraction of signaling proteins such as Cyt C from cytoplasma directly without the need of isolation of subcellular organelles such as mitochondria. Also, it does not require cell lysis or the addition of any reagents into the cells, minimizing the variability and increasing the assay robustness. As such, this approach is very time-efficient, taking only 15 min totally, as compared with hours and even days by Western blotting. Third, the current approach is ultrasensitive, allowing for the detection of signaling proteins at the single molecule level. This enabled the monitoring of signaling proteins at the early events of a signaling pathway. In addition to apoptosis, the scPISA approach can be applied to many other signaling pathways.

We further demonstrated the scPISA approach as a promising tool for precision assessing anti-cancer efficacy of cytotoxic drugs. We proposed two key signaling proteins in the apoptosis pathway, including Cyt C and caspase-3, as apoptosis indicators. In terms of the expression level of these apoptosis indicators, we proposed a set of new parameters, including *I*_max_, *T*_50_ and *EC*_50_, for the comprehensive and quantitative evaluation of the efficacy of anti-cancer drugs. As compared with conventional assays for drug efficacy evaluation such as MTT, this approach showed several unique strengths. First, the scPISA requires only a limited number of cells, ca. 60 cells for one whole set of assays. As a comparison, conventional assays need a much larger population of cells. Thus, this approach holds incomparable potential for personalized drug screening and precision medicine. Because a patient’s cancer cells from biopsies or surgically resected tumors are usually limited whereas cultured cancer cells are not representative of those in the patient, conventional assays may fail to reflect the drug sensitivity of the patient’s cancer cells. However, the scPISA approach permits directly measuring the cells from biopsies or surgically resected tumors from the patient, without the need of further culture. Second, the scPISA approach is signaling pathway-specific, providing important insights into the molecular mechanism underlying cancer killing. Third, this approach is more reliable. It relies on two correlated signaling proteins, one upstream and one downstream, as well as a comprehensive set of quantitative evaluation parameters. The two apoptosis indicators are mutually checkable and so are the three parameters, providing an error-proofing mechanism. Last, the scPISA approach is much more sensitive than conventional assays particularly MTT. When a large population of cells is used, some died cells or dying cells are very likely involved in. This may give over-estimated results. As a contrast, as single living cells are used in scPISA approach, such possibility can be avoided and thereby the results should be more reliable. Due to these significant merits, the scPISA approach can be a promising tool for facile anti-cancer efficacy evaluation in precision medicine and personalized treatment of cancer.

## ONLINE METHODS

### scPISA of signaling proteins and protein-protein complexes

The principle and procedure of scPISA is illustrated in **Scheme 1**. The procedure included three major steps: 1) in-cell extraction, 2) labeling, and 3) detection. To probe Cyt C or caspase-3, an extraction probe immobilized with anti-Cyt C or anti-caspase-3 antibody was precisely inserted into a single apoptotic cell under investigation through an in-lab built three-dimensional cell manipulation platform. After insertion, the probe was kept in the cell for 5 min to extract target signaling proteins. Then, the probe was taken out of the cell and washed with 100 mM phosphate buffer, pH 7.4, for three times, target protein molecules captured by the extraction probe were labeled by dipping with 5 µL of anti-Cyt C or anti-caspase-3 antibody immobilized PATP-encapsulated Ag@SiO_2_ NPs for 5 min. Then the extraction probe was washed with 100 mM phosphate buffer, pH 7.4, for three times, dried and then detected by the Raman spectrograph. For the Raman detection, map scanning mode was used, scanning an area of 2 × 5 μm (width × length) on the probe line-by-line for 4 times (0.5 µm width and 10 spots for each line) starting from about 1 μm away from the tip. Among the obtained 40 Raman spectra, spectra without Raman signal were discarded and the averaged intensity for the middle 6 spots of spectra with Raman signal was taken for scPISA assay. To probe survivin associated protein-protein complexes, an anti-Cyt C or anti-caspase-3 antibody-immobilized microprobe was used for in-cell extraction whereas anti-survivin polyclonal antibody-immobilized PATP-encapsulated Ag@SiO_2_ NPs were used for labeling. The other procedure was the same as above.

### Evaluation of anti-cancer drugs-induced apoptosis by scPISA

To measure the whole set of apoptosis evaluation parameters, experiments on time-response relationship and dose-response dependence were performed. To establish the time-response relationship, cells under test were treated with same apoptotic stimuli for different time. After a stimulus of a certain time, apoptosis indicator proteins were detected by scPISA. Average Raman intensity for 3 individual cells was plotted against stimulation time. Logistic function was used to fit these plots, giving the values for *T*_50_ and *I*_max_. To develop the dose-response dependence, cells under test were stimulated with an anti-cancer drug alone of different concentrations or its combination with another drug for an appropriate simulation time. Then, apoptosis indicator proteins were detected by scPISA. Using the average Raman intensity for 3 individual cells for each concentration point, dose-response dependences were drawn. Fitting of the dose-dependent response curve with the logistic function yielded values for the parameters *EC*_50_ and *I*_max_.

## Supporting information

Supplementary file for Probing apoptosis signaling proteins in single living cells for precision efficacy evaluation of anti-cancer drugs

## ACKNOWLEDGMENTS

We acknowledge financial support of the Key Scientific Instrumentation Grant (21627810) and the National Science Fund for Distinguished Young Scholars (21425520) from the National Natural Science Foundation of China, and Excellent Research Program of Nanjing University (ZYJH004).

## AUTHOR CONTRIBUTIONS

Z.L. conceived the study and designed the experiments. Y.R.W. performed most of the experiments and analyzed the data. J.L. provided intellectual guidance in the design of partial experiments. H.H. performed some experiments on characterization and verification. Z.L. and Y.R.W. wrote the paper.

## References

1. Nault, J.C., Galle, P.R. & Marquardt, J.U. The role of molecular enrichment on future therapies in hepatocellular carcinoma. J. Hepatol. 69, 237–247 (2018).

2. Wehling, M. Assessing the translatability of drug projects: what needs to be scored to predict success? Nat. Rev. Drug Discov. 8, 541–546 (2009).

3. Sharma, S.V., Haber, D.A. & Settleman, J. Cell line-based platforms to evaluate the therapeutic efficacy of candidate anticancer agents. Nat. Rev. Cancer 10, 241–253 (2010).

4. Barretina, J. et al. The Cancer Cell Line Encyclopedia enables predictive modelling of anticancer drug sensitivity. Nature 483, 603–607 (2012).

5. Fallahi-Sichani, M., Honarnejad, S., Heiser, L.M., Gray, J.W. & Sorger, P.K. Metrics other than potency reveal systematic variation in responses to cancer drugs. Nat. Chem. Biol. 9, 708–714 (2013).

6. Alberts, D.S. et al. In-vitro clonogenic assay for predicting response of ovarian cancer to chemotherapy. Lancet 2, 340–342 (1980).

7. Yin, T. et al. Combined effects of As4S4 and imatinib on chronic myeloid leukemia cells and BCR-ABL oncoprotein. Blood 104, 4219–4225 (2004).

8. Feng, Y., Mitchison, T.J., Bender, A., Young, D.W. & Tallarico, J.A. Multi-parameter phenotypic profiling: using cellular effects to characterize small-molecule compounds. Nat. Rev. Drug Discov. 8, 567–578 (2009).

9. Friedman, A.A., Letai, A., Fisher, D.E. & Flaherty, K.T. Precision medicine for cancer with nextgeneration functional diagnostics. Nat. Rev. Cancer 15, 747–756 (2015).

10. Andersen, J.N. et al. Pathway-Based Identification of Biomarkers for Targeted Therapeutics: Personalized Oncology with PI3K Pathway Inhibitors. Sci. Transl. Med. 2, 14 (2010).

11. Sevecka, M. & MacBeath, G. State-based discovery: a multidimensional screen for small-molecule modulators of EGF signaling. Nat. Methods 3, 825–831 (2006).

12. Conway, J.R., Carragher, N.O. & Timpson, P. Developments in preclinical cancer imaging: innovating the discovery of therapeutics. Nat. Rev. Cancer 14, 314–328 (2014).

13. Pemovska, T. et al. Individualized Systems Medicine Strategy to Tailor Treatments for Patients with Chemorefractory Acute Myeloid Leukemia. Cancer Discov. 3, 1416–1429 (2013).

14. Altschuler, S.J. & Wu, L.F. Cellular Heterogeneity: Do Differences Make a Difference? Cell 141, 559–563 (2010).

15. Anchang, B. et al. DRUG-NEM: Optimizing drug combinations using single-cell perturbation response to account for intratumoral heterogeneity. P. Natl. Acad. Sci. USA 115, E4294–e4303 (2018).

16. Krutzik, P.O., Crane, J.M., Clutter, M.R. & Nolan, G.P. High-content single-cell drug screening with phosphospecific flow cytometry. Nat. Chem. Biol. 4, 132–142 (2008).

17. Ullal, A.V. et al. Cancer Cell Profiling by Barcoding Allows Multiplexed Protein Analysis in Fine-Needle Aspirates. Sci. Transl. Med. 6, 11 (2014).

18. Austin, L.A., Kang, B. & El-Sayed, M.A. A New Nanotechnology Technique for Determining Drug Efficacy Using Targeted Plasmonically Enhanced Single Cell Imaging Spectroscopy. J. Am. Chem. Soc. 135, 4688–4691 (2013).

19. Bendall, S.C. et al. Single-cell mass cytometry of differential immune and drug responses across a human hematopoietic continuum. Science 332, 687–696 (2011).

20. Gaudet, S. & Miller-Jensen, K. Redefining Signaling Pathways with an Expanding Single-Cell Toolbox. Trends Biotechnol. 34, 458–469 (2016).

21. Hughes, A.J. et al. Single-cell western blotting. Nat. methods 11, 749 (2014).

22. Miller, M.A. & Weissleder, R. Imaging of anticancer drug action in single cells. Nat. Rev. Cancer 17, 399–414 (2017).

23. Shi, T.J. et al. Conservation of protein abundance patterns reveals the regulatory architecture of the EGFR-MAPK pathway. Sci. Signal. 9, 13 (2016).

24. Adlung, L. et al. Protein abundance of AKT and ERK pathway components governs cell type-specific regulation of proliferation. Mol. Syst. Boil. 13, 904 (2017).

25. Liu, J., Yin, D.Y., Wang, S.S., Chen, H.Y. & Liu, Z. Probing Low-Copy-Number Proteins in a Single Living Cell. Angew. Chem. Int. Edit. 55, 13215–13218 (2016).

26. Tu, X.Y. et al. Molecularly Imprinted Polymer-Based Plasmonic Immunosandwich Assay for Fast and Ultrasensitive Determination of Trace Glycoproteins in Complex Samples. Anal. Chem. 88, 12363–12370 (2016).

27. Li, W. et al. Controllably Prepared Aptamer-Molecularly Imprinted Polymer Hybrid for High-Specificity and High-Affinity Recognition of Target Proteins. Anal. Chem. 91, 4831–4837 (2019).

28. Spencer, S.L. & Sorger, P.K. Measuring and modeling apoptosis in single cells. Cell 144, 926–939 (2011).

29. Green, D.R. Apoptotic pathways: Ten minutes to dead. Cell 121, 671–674 (2005).

30. Garrido, C. et al. Mechanisms of cytochrome c release from mitochondria. Cell Death Differ. 13, 1423–1433 (2006).

31. Riedl, S.J. & Shi, Y.G. Molecular mechanisms of caspase regulation during apoptosis. Nat. Rev. Mol. Cell Bio. 5, 897–907 (2004).

32. Julien, O. & Wells, J.A. Caspases and their substrates. Cell Death Differ. 24, 1380–1389 (2017).

33. Gray, D.C., Mahrus, S. & Wells, J.A. Activation of Specific Apoptotic Caspases with an Engineered Small-Molecule-Activated Protease. Cell 142, 637–646 (2010).

34. Li, P. et al. Cytochrome c and dATP-dependent formation of Apaf-1/caspase-9 complex initiates an apoptotic protease cascade. Cell 91, 479–489 (1997).

35. Igney, F.H. & Krammer, P.H. Death and anti-death: Tumour resistance to apoptosis. Nat. Rev. Cancer 2, 277–288 (2002).

36. Shaffer, S.M. et al. Rare cell variability and drug-induced reprogramming as a mode of cancer drug resistance. Nature 546, 431–435 (2017).

37. Tamm, I. et al. IAP-family protein Survivin inhibits caspase activity and apoptosis induced by Fas (CD95), Bax, caspases, and anticancer drugs. Cancer Res. 58, 5315–5320 (1998).

38. Deveraux, Q.L. & Reed, J.C. IAP family proteins--suppressors of apoptosis. Gene Dev. 13, 239–252 (1999).

39. Dohi, T., Beltrami, E., Wall, N.R., Plescia, J. & Altieri, D.C. Mitochondrial survivin inhibits apoptosis and promotes tumorigenesis. J. Clin. Invest. 114, 1117–1127 (2004).

40. Singh, N., Krishnakumar, S., Kanwar, R.K., Cheung, C.H.A. & Kanwar, J.R. Clinical aspects for survivin: a crucial molecule for targeting drug-resistant cancers. Drug Discov. Today 20, 578–587 (2015).

41. Liu, Z., Delavan, B., Roberts, R. & Tong, W. Lessons Learned from Two Decades of Anticancer Drugs. Trends Pharmacol. Sci. 38, 852–872 (2017).

42. Arunachalam, G., Samuel, S.M., Marei, I., Ding, H. & Triggle, C.R. Metformin modulates hyperglycaemia-induced endothelial senescence and apoptosis through SIRT1. Brit. J. Pharmacol. 171, 523–535 (2014).

43. Wu, L. et al. An Ancient, Unified Mechanism for Metformin Growth Inhibition in C. elegans and Cancer. Cell 167, 1705–1718.e1713 (2016).

