## Supplementary file for Probing apoptosis signaling proteins in single living cells for precision efficacy evaluation of anti-cancer drugs for "Probing apoptosis signaling proteins in single living cells for precision efficacy evaluation of anti-cancer drugs"

### EXPERIMENTAL SECTION

#### 1. Reagents and Materials

3-Aminopropyltriethoxysilane (APTES), dopamine hydrochloride, tetraethoxysilane (TEOS), 4-aminothiophenol (PATP), bovine serum albumin (BSA), chloroauric acid ($HAuCl_4$), silver nitrate ($AgNO_3$), glacial acetic acid (HAc), sodium cyanoborohydride, HCl (36%), trisodium citrate, NaOH, $NaH_2PO_4$, $Na_2HPO_4$, $KHCO_3$, methanol and anhydrous ethanol were purchased from Nanjing Reagent Company (Nanjing, China). Polyclonal anti-survivin antibody (ab24479), monoclonal anti-survivin antibody (ab134170), monoclonal anti-active caspase-3 antibody (ab32042) polyclonal anti-active caspase-3 antibody (ab2302), and recombinant human active caspase-3 full length protein (ab52314) were purchased from Abcam (Shanghai, China). Recombinant human Cytochome C (Cyt C) full length protein (230-00671-50) was purchased from Raybiotech (Shanghai, China). Actinomycin D (Act D) was purchased from Solarbio Life Sciences (Shanghai, China). Metformin was obtained from Sigma-Aldrich (Shanghai, China). Monoclonal anti-Cyt C antibody (12963s) was perchased from Cell Signaling Technology (Shanghai, China). Polyclonal anti-Cyt C antibody (ARG20006) was perchased from Arigo (Shanghai, China). Sodium dodecyl sulfate (SDS) was obtained from Bio-Rad (Hercules, CA, USA). Staurosporine (STS), 11-mercaptoundecanoic acid (11-MUA), 1-ethyl-3-(3'-dimethylaminopropyl) carbodiimide (EDC), *N*-hydroxysuccinimide (NHS), and 4-(2-hydroxyethyl)-1-piperazineethanesulfonic acid (HEPES) were from J&K Scientific (Shanghai, China). Normal hepatic cells (L-02), human hepatoma carcinoma cells (HepG-2), human cervical cancer cells (HeLa), Dulbecco's modified eagle medium (DMEM, containing 4.5 mg/mL glucose, 80 U/mL penicillin and 0.08 mg/mL streptomycin), 1x phosphate-buffered saline (PBS) solution and MTT Cell Proliferation and Cytotoxicity Assay Kit were from Keygen Biotech (Nanjing, China). Water used in all experiments was purified with a Milli-Q Advantage A10 system (Millipore, Milford, MA, USA). Other chemicals were all of ananlytical purity without further treatment. Borosilicate glass capillaries of 0.58 mm i.d. and 1.0 mm o.d. were purchased from Naturegene Life Sciences (Shanghai, China).

### 2. Instrumentation

Plasmonic detection was carried out on a Renishaw InVia Reflex confocal microscope (Renishaw, UK) equipped with a high-resolution grating with 1,800 grooves/mm, additional band-pass filter optics, and a CCD camera. Spectra were acquired using a 633 nm excitation laser line (1 s integration time and 1 accumulation). The laser was focused onto the sample by using a ×50 objective (N.A. 0.75), providing a spatial resolution of ca. 1 $\mu m^2$. All measurements were carried out using a He-Ne laser ($\lambda_0$ = 633 nm; laser power at spot, ca. 8.5 mW). Wavelength calibration was performed by measuring silicon wafers though a ×50 objective, evaluating the first-order phonon band of Si at 520 $cm^{-1}$. Each spectrum was baseline corrected except noise test. The microprobes used in this work were fabricated by tapering 1.0 mm core-diameter borosilicate glass capillaries using a P-2000 F pipette puller (Sutter Instrument, Novato, CA, USA), producing probes with a diameter of about 700 nm. An in-lab built three-dimensional cell manipulation platform, composed of a TransferMan® 4r micromanipulator (Eppendorf, Germany), an InjectMan® 4 micromanipulator and a PiezoXpert® piezo-assisted cell membrane penetrator, mounted on an inverted microscope (Olympus IX73, Japan), was used to precisely insert extraction microprobes into single cells under investigation. Scanning electron microscopic (SEM) characterization was performed on a FE-SEM S-4800 instrument (Hitachi, Tokyo, Japan). Transmission electron microscopic (TEM) characterization was carried out on a JEM-1011 system (JEOL, Tokyo, Japan).The UV-VIS was obtained from MTT assay was based on the signal read from a Synergy Mx microplate reader (BioTek, Winooski, VT, USA).

### 3. Preparation of Extraction Microprobes

The preparation procedure of the probe is illustrated in **Supplementary Fig. 1a**. First, Au-coated microprobes were prepared as supporting microprobes. Then, the supporting microprobes were modified with antibody. The detailed procedures are described below.

#### 3.1. Preparation of Au-coated Supporting Microprobes

A gold layer was prepared onto the surface of supporting microprobes according to the chemical plating method.[1] Briefly, the probes were immersed in a mixed solution (12 mM $HAuCl_4$, 0.5

M $KHCO_3$ and 25 mM glucose) for 3-4 h at 50 °C (air bath) until an obvious gold layer appeared on the surface of each probe. Then, the Au-coated probes were washed with water and then dried by air at room temperature.

#### 3.2. Preparation of Antibody-immobilized Microprobes

The procedure included three major steps: carboxy-functionalization, antibody immobilization and BSA sealing. For carboxy-functionalization, Au-coated supporting microprobes were first immersed in 11-mercaptoundecanoic acid (1 mg/mL, dissolved in ethanol) at room temperature for 12 h, followed by rinse with ethanol for three times, and dried. To immobilize antibody onto the prepared probes, the probes were immersed in 1 mL of 1:1 EDC: NHS (100 and 25 mM, respectively) for 10 min. The probes were washed twice with phosphate solution (100 mM, pH 4.5), and dried. They were then immersed in 1 mL of 10 μg/mL desired antibody and allowed for reaction for 2 h. To reduce non-specific adsorption of non-target proteins onto the probes, the probes were sealed with BSA. To do that, the probes were first washed twice with 100 mM phosphate buffer (pH 7.4), and then immersed in 1 mL of 2.5% BSA in 100 mM phosphate buffer (pH 7.4) for 2 h. Finally, the antibody-modified probes were washed with 100 mM phosphate buffer (pH 7.4) for three times and dried at room temperature, then stored at 4 °C for further use.

### 4. Preparation of Raman Nanotags

AgNPs were first prepared as base nanotags. Then, Raman-active AgNPs were prepared according to the preparation procedures shown in **Supplementary Fig. 1b**. The detailed procedures are described below.

#### 4.1. Synthesis of Silver Nanoparticles (AgNPs)

AgNPs were prepared as described by Lee and Meisel.[2] Briefly, $AgNO_3$ (36 mg) was dissolved in 200 mL water and brought to boil under continuous stirring. Then, 4 mL of 1% (w/v) trisodium citrate was added. The mixture was boiled with stirring for about 1 h and then cooled down to room temperature. The obtained AgNPs solution was stored at 4 °C for further use.

### 4.2. Fabrication of Raman Nanotags for Antibody-immobilized Microprobes

PATP-encapsulated antibody-modified AgNPs were prepared and used as Raman nanotags. The procedure involved in four steps, including modification with PATP, encapsulation of PATP-AgNPs, functionalization with APTES, and antibody immobilization.

To modify AgNPs with PATP, 20 μL of 1 mM PATP (dissolved in ethanol) was added to 10 mL AgNPs stock solution and the mixed solution was stirred for 1 h at room temperature. To encapsule PATP-AgNPs with a silica shell, 40 mL of ethanol, 700 μL ammonia and 10 mL of 10 mM TEOS (dissolved in ethanol) were added with stirring for about 70 min. Centrifugation was performed at 10,000 rpm for 10 min. Then the particles were resuspended in 50 mL ethanol. To functionalize the PATP-encapsuled AgNPs with APTES, 150 μL of APTES was added and stirred for 1 h. Then the amino-modified AgNPs were separated from solution by centrifugation at 10,000 rpm for 10 min. The clear supernatant was discarded, and the loosely packed silver sediment was collected. To immobilize the amino-modified AgNPs with antibody, the collected sediment was first resuspended in 10 mL glutaraldehyde solution (0.5%) with stirring for 2 h, followed by rinse with ultrapure water for three times, then resuspended in 10 mM HEPES solution. After that, the Raman reporter-encapsulated AgNPs were immobilized with anti-Cyt C or anti-caspase-3 or anti-survivin polyclonal antibody. While gently agitating, 10 μg/mL of antibody was added to 10 mL suspension of the reporter-encapsulated silver colloids with stirring for 2 h at room temperature. Upon centrifugation at 10,000 rpm for 10 min, two phases were obtained: a clear supernatant of unbound antibody and a loosely packed sediment of the reporter-encapsulated antibody-immobilized AgNPs. The obtained NPs were washed with 100 mM phosphate buffer (pH 7.4) for 5 min. Finally, the antibody-immobilized Raman-active AgNPs were dispersed in a 100 mM phosphate buffer (pH 7.4) and stored at 4 °C for further use.

### 5. Dependence of Raman intensity on signaling protein concentration

Each antibody-immobilized microprobe was dipped with a 5-μL solution of target protein (Cyt

C or caspase-3) of different concentrations, dissolved in 100 mM phosphate buffer (pH 7.4) for 20 min. After washing with 100 mM phosphate buffer (pH 7.4) for three times, captured protein was labeled with 10 μL of Raman-active antibody-immobilized AgNPs for 5 min. Then the microprobe was rinsed with 100 mM phosphate buffer (pH 7.4) for three times, dried and then detected by the Raman microscope.

### 6. Cell Viability Test and $IC_{50}$ Calculation

Cell viability was measured by the MTT assay. Briefly, Hela cells were seeded at the density of $2\times10^4$ cells/well in 96-well plate. Cells were treated with Act D (4 μM) alone or metformin alone or in their combination for 24 h . Then MTT (5 mg/mL) was added and incubated for 4 h at 37 °C. After that, the medium with MTT was removed out, and 100 μL of DMSO was added to each well, and the 96-well plate was left at room temperature in the dark and shaken well with a shaker for 2 hours. Absorbance at 490 nm in each well, including the blanks, were measured in a microplate reader. The cell viability was calculated according to the equation.

$$Cell\ viability\ \% = \left(OD_{tested\ drug}/OD_{control}\right) \times\ 100\ \% \qquad (1)$$

The $IC_{50}$ value of tested drug was obtained by fitting the data according to the following equation.

$$Y = Min + \frac{Max-Min}{1+(\frac{X}{IC_{50}})^h} \qquad (2)$$

*X* is the logarithm of drug concentration, *Y* is the Cell viability %, and *h* is the Hill coefficient. *Min* stands for the minimum *Y* value, while *Max* stands for the maximal *Y* value.

### 7. scPISA of Signaling Proteins in Single Cells

An extraction probe immobilized with anti-Cyt C or anti-caspase-3 antibody was precisely inserted into a single apoptotic cell through the three-dimensional manipulator. After insertion, the probe was kept in the cell for 5 min to extract target signaling proteins. Then, the probe was taken out from the cell and washed with 100 mM phosphate buffer, pH 7.4, for three times, target protein molecules captured by the extraction probe were labeled by dipping with 5 μL of anti-Cyt C or anti-caspase-3 antibody immobilized AgNPs for 5 min. Then the probes were washed with 100 mM phosphate buffer, pH 7.4, for three times, dried and then detected by the Raman spectrograph.

Sister cells were choosed to examine the reliability of scPISA for probing signaling proteins. We tracked the HepG-2 cell lines for 7 hours under microscopy after the apoptosis inducers were added, 2 new divided daughter cells from a mother cell, namely, a group of sister cells were choosed for signaling ptorein analysis. The microprobe was inserted into the cells approximately with the same depth of 5-6 μm, in a fixed angle of 45 degree, and was kept 1-2 μm away from the nucleus, under the microscope. Then it followed the same procedures above.

### References

1. Zhou, F. et al. Sensitive sandwich ELISA based on a gold nanoparticle layer for cancer detection. *Analyst* **137**, 1779-1784 (2012).
2. Lee, P. C. , and D. J. J. Meisel . Adsorption and Surface-Enhanced Raman of Dyes on Silver and God Sols. *J. Phy. Chem.* **86**, 3391-3395 (1982).

### SUPPLEMENTARY DATA

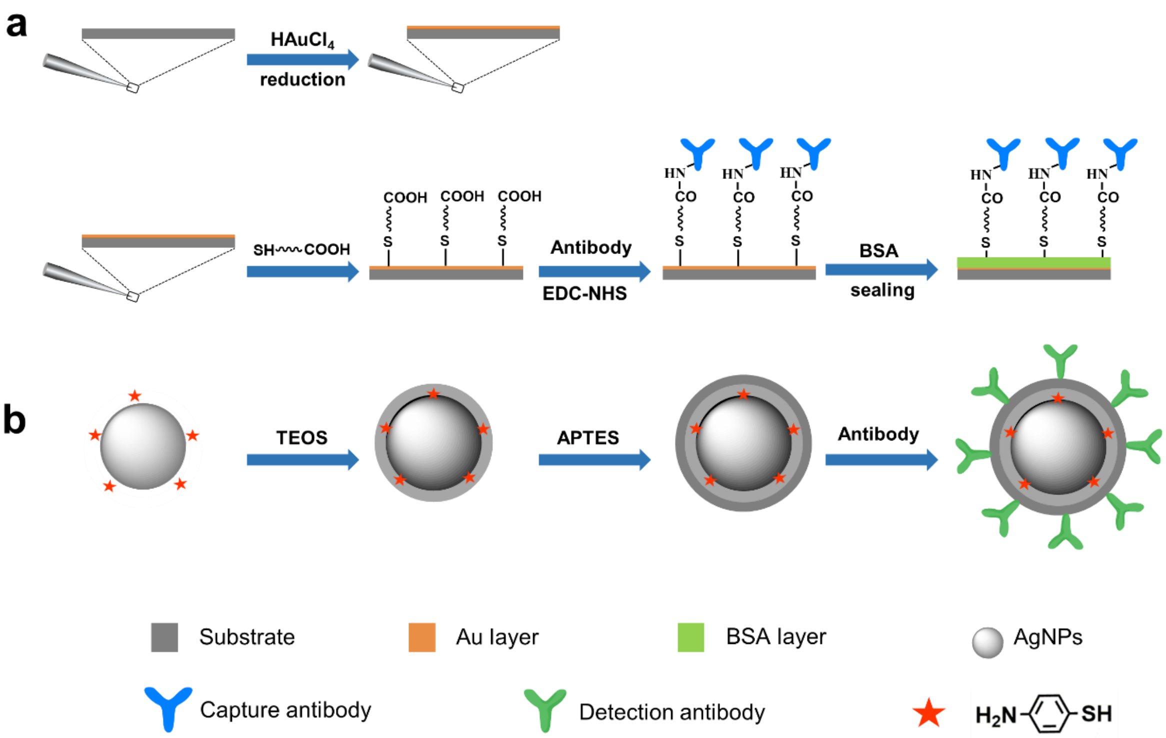


**Supplementary Figure 1. Prodedure for the preparation of extraction microprobes (a) and Raman nanotags (b).**

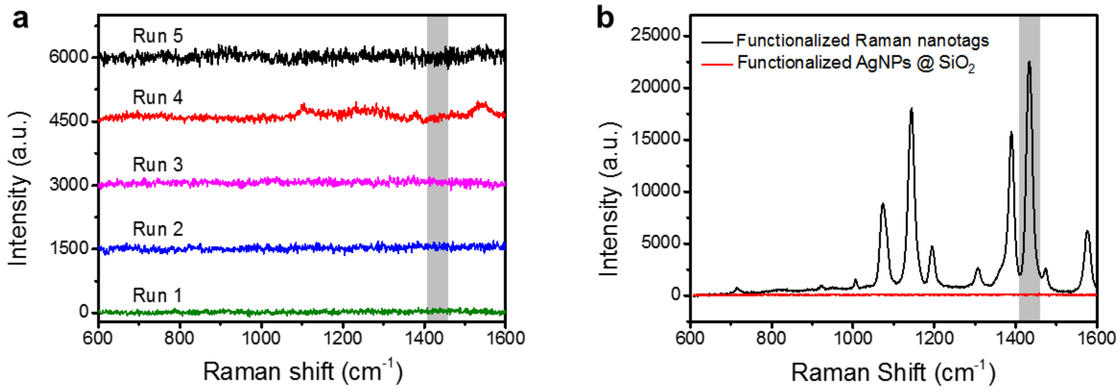


**Supplementary Figure 2. Characterization of extraction microprobe (a) and Raman nanotags (b).**

**(a)** Raman spectra for anti-caspase-3 monoclonal antibody-functionalized microprobe. **(b)** Raman spectra for anti-caspase-3 antibody-functionalized Raman nanotags (peak indicated by grey band was used for the scPISA assay) and anti-caspase-3 antibody-functionalized AgNPs@$SiO_2$ (without Raman reporter).

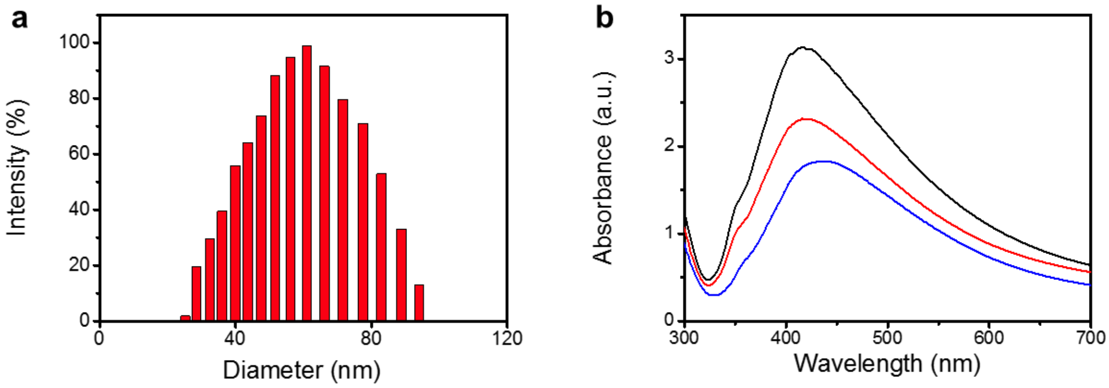


**Supplementary Figure 3. Characterization of Raman nanotags.**

**(a)** DLS of AgNPs. **(b)** UV-vis absorption spectra of AgNPs (black), PATP-encapsulated Ag@$SiO_2$ NPs (red) and antibody-functionalized PATP-encapsulated Ag@$SiO_2$ NPs (final Raman nanotags) (blue).

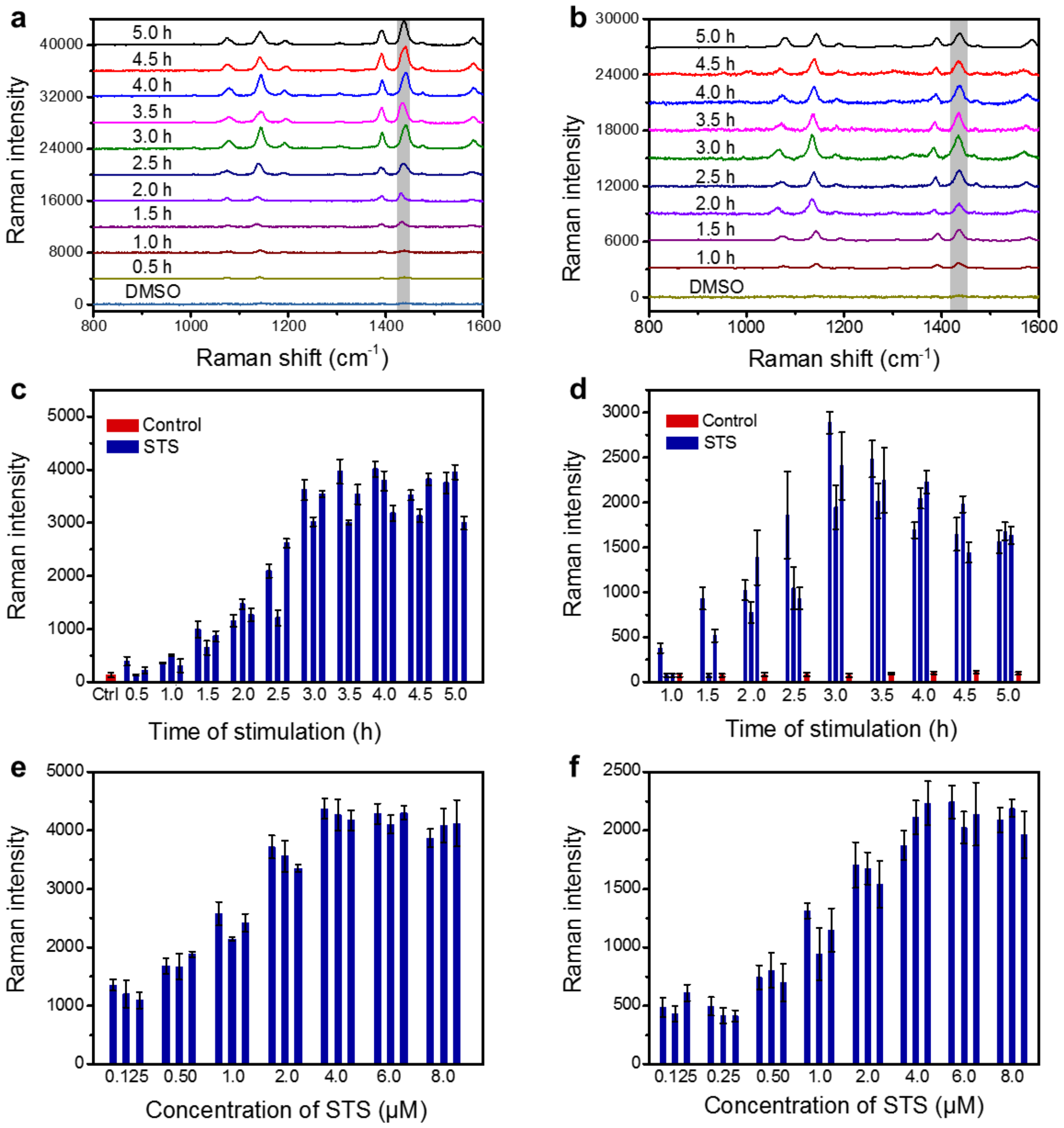


**Supplementary Figure 4. Probing signaling proteins in apoptotic single cells.**

Representative Raman spectra for **(a)** Cyt C and **(b)** caspase-3 in single apoptotic Hela cells stimulated with 4 μM STS for different time. Raman intensity for **(c)** Cyt C and **(d)** caspase-3 in single Hela cells stimulated with 4 μM STS for different time. Error bars represent standard deviations for Raman signals from 6 spots on a single microprobe. Raman intensity for **(e)** Cyt C and **(f)** caspase-3 in single Hela cells stimulated with STS of different concentration for 3 h. Cotrols were cells incubated in the culture medium containing 1‰ (v/v) DMSO.

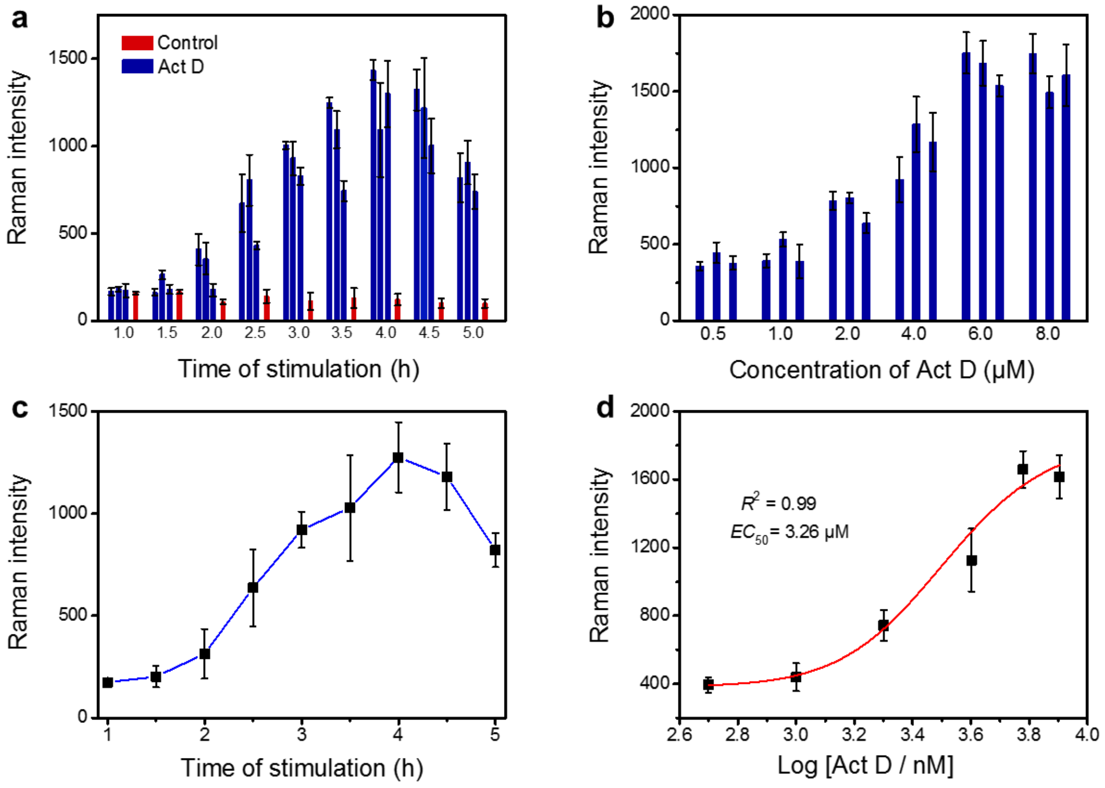


**Supplementary Figure 5. Raman intensity for caspase-3 in apoptotic Hela cells induced by Act D**.

**(a)** Raman intensity for caspase-3 in single Hela cells stimulated with 4 μM Act D for different time. Error bars represent standard deviations for Raman signals from 6 spots on a single microprobe. **(b)** Raman intensity for caspase-3 in single Hela cells stimulated with Act D of different concentration for 4 h. **(c)** Dependence of the average Raman intensity for caspase-3 in single Hela cells stimulated with 4 μM Act D on the stimulation time. Error bars represent standard deviations for Raman signals from 3 single cells. **(d)** Dose-effect curve with caspase-3 as the indicator for the apoptosis of Hela cells induced by Act D.

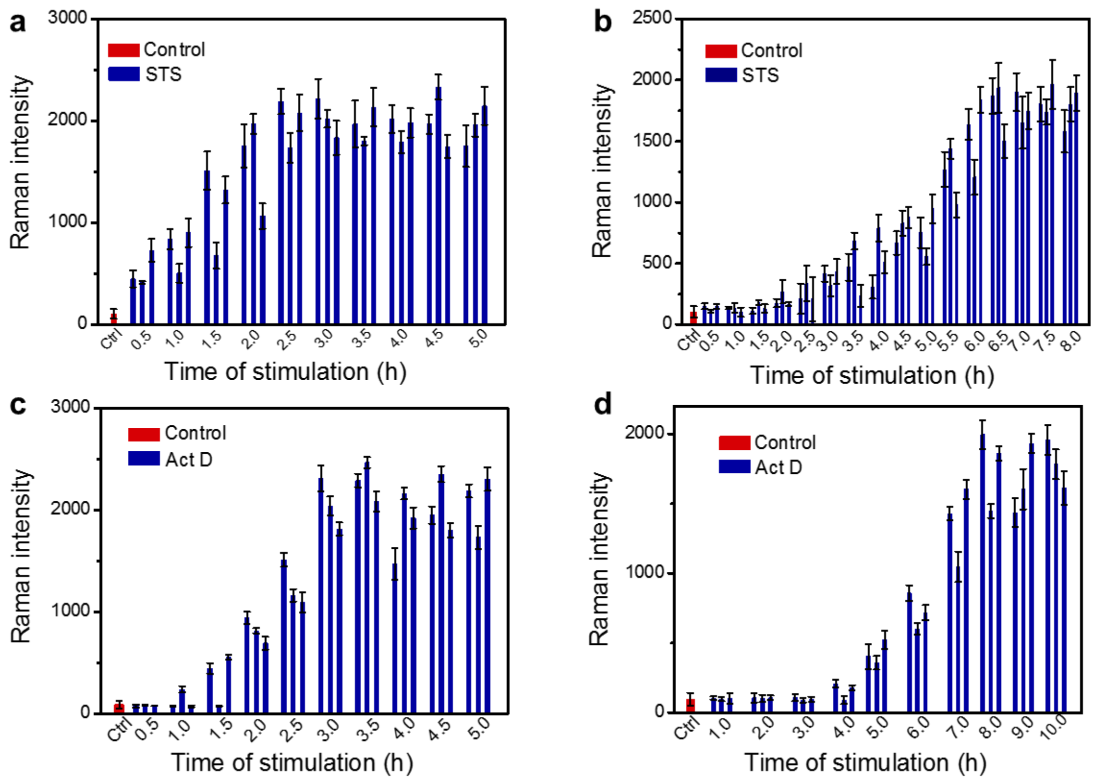


**Supplementary Figure 6. Comparison of the time-dependent expression level of Cyt C in single apoptotic normal hepatic cells (L-02) and hepatoma carcinoma cells (HepG-2).** Raman intensity for Cyt C in **(a)** L-02 cells and **(b)** HepG-2 cells stimulated with 4 μM STS for different time, and Raman intensity for Cyt C in **(c)** L-02 cells and **(d)** HepG-2 cells stimulated with 4 μM Act D for different time. Error bars represent standard deviations for Raman signals from 6 spots on a single microprobe. Controls were cells incubated in the culture medium containing 1‰ (v/v) DMSO, the Raman intensity for the controls was the average intensity for the control experiments at all the time.

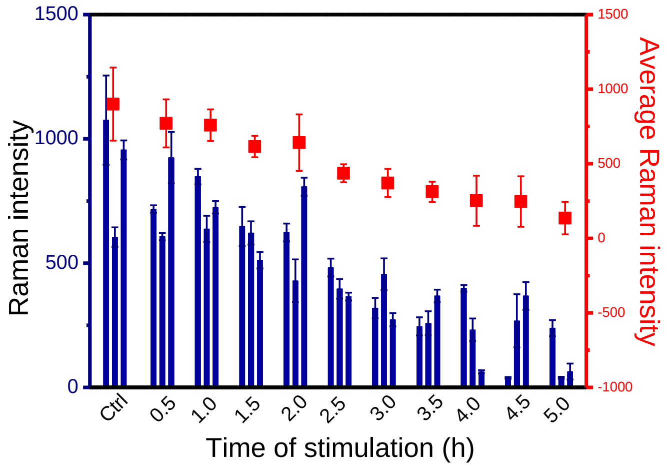


**Supplementary Figure 7. Decay of the expression level of survivin in apoptotic HepG-2 cells along the stimulation time.**

Raman intensity for survivin in single HepG-2 cells stimulated with 4 μM STS for different time (blue). Average Raman intensity for survivin in 3 single cells (red). Error bars in blue represent standard deviations for Raman signals from 6 spots on a single microprobe, while error bars in red represent standard deviations for Raman signals from 6 spots on a single microprobe.

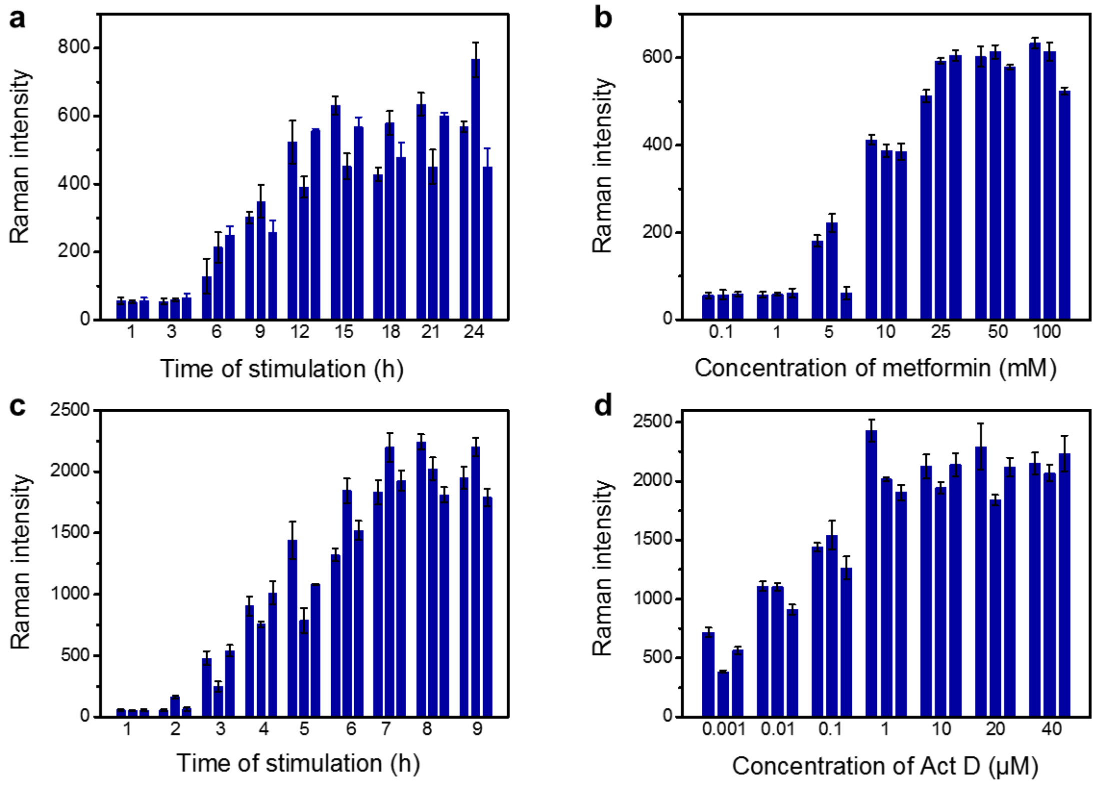


**Supplementary Figure 8. Evaluation of apoptosis inducement efficacy of metformin used alone (a and b) and in combination with Act D (c and d) by scPISA.**

Raman intensity for Cyt C in single HepG-2 cells stimulated with 10 mM of metformin **(a)** and in combination with Act D (4 μM) **(c)** for different time. Raman intensity for Cyt C in single HepG-2 cells stimulated by metformin of defferent concentration for 12 h **(b)** and by metformin (10 mM) in combination with Act D of defferent concentration for 8 h **(d)**. Error bars represent standard deviations for Raman signals from 6 spots on a single microprobe.

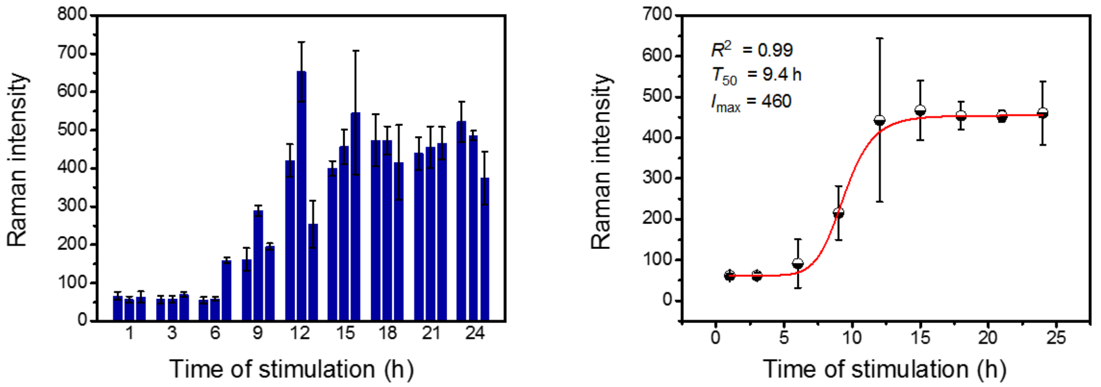


**Supplementary Figure 9. Expression level of caspase-3 in single apoptostic cells induced by metfoemin.**

**(a)** Raman intensity for caspase-3 in single HepG-2 cells stimulated with 10 mM of metformin for different time. **(b)** Logistic fitting of the average Raman intensity for caspase-3 in apoptotic HepG-2 cells stimulated with 10 mM of metformin for different time.

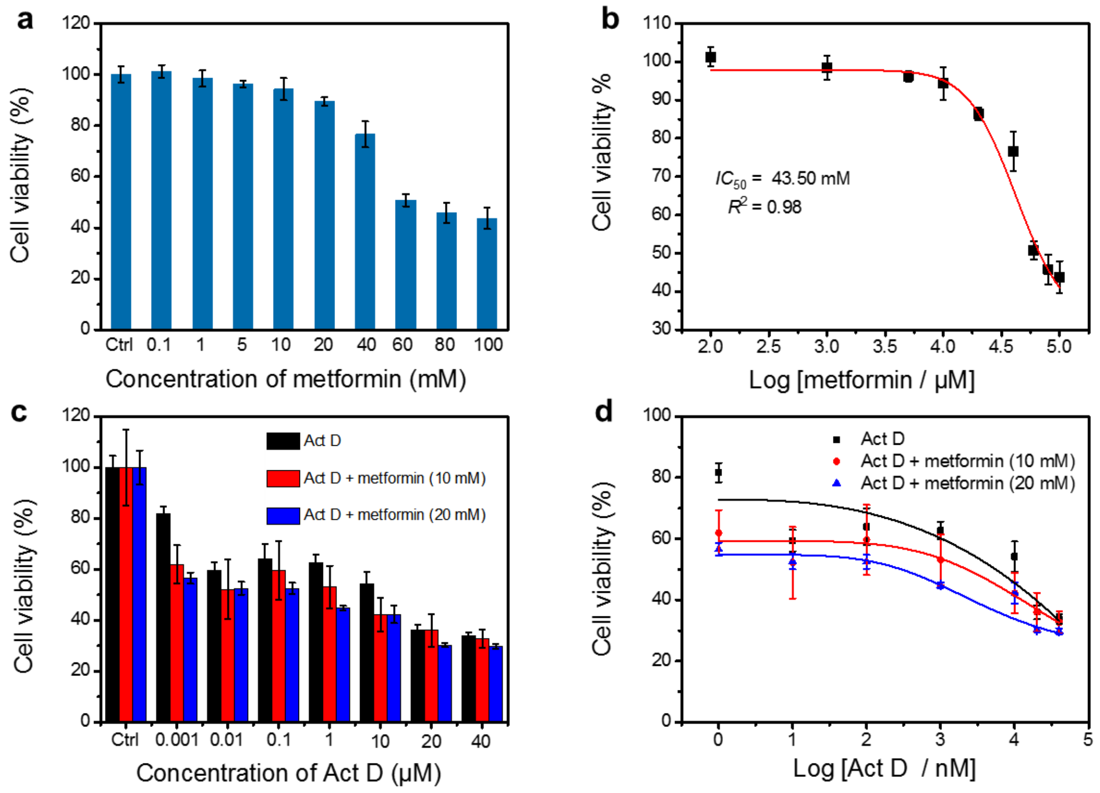


**Supplementary Figure 10. MTT evaluation of the anti-cancer efficacy of metformin alone (a and b) and in combination with Act D of different concentration (c and d).**

**(a)** Cell viability of HepG-2 cells after incubated with metformin alone of different concentration for 24 h and **(b)** calculation of $IC_{50}$ using data given in **a**. **(c)** Cell viability of HepG-2 cells after incubated with Act D of different concentrations without and with 10 mM or 20 mM metformin for 24 h. **(d)** Calculation of $IC_{50}$ using data given in **c**.

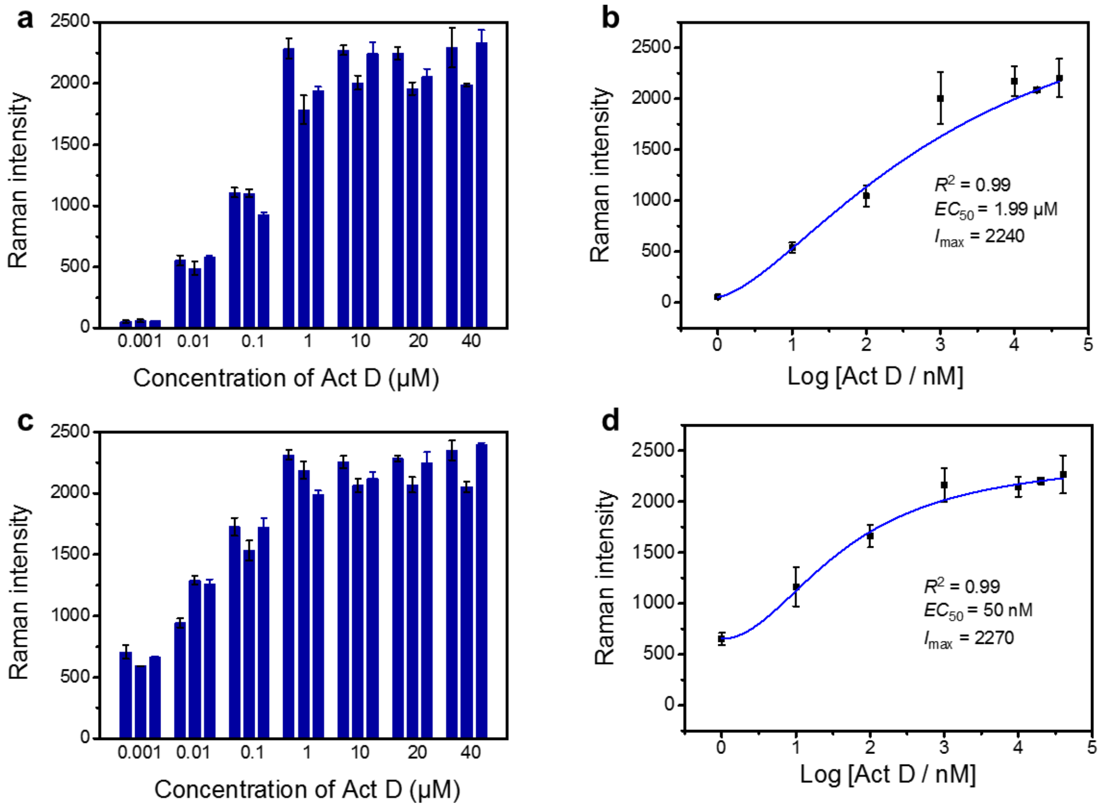


**Supplementary Figure 11. scPISA evaluation of the anti-cancer efficacy of Act D alone (a and b) in combination with 20 mM metformin (c and d).**

Raman intensity for Cyt C in single HepG-2 cells co-stimulated with Act D (4μM) alone **(a)** and in combination with metformin (20 mM) **(c)** for 12 h. Error bars represent standard deviations for Raman signals from 6 spots on a single microprobe. Dose-effect curve for apoptosis induced by Act D alone **(b)** and in combination with metformin (20 mM) **(d)** for 12 h, using Cyt C as an apoptosis indicator. Error bars represent standard deviations for Raman signals from 3 single cells.

**Supplementary Table 1. Comparison of the anti-cancer efficacyof STS and Act D towards Hela cells with caspase-3 as the indicator of apoptosis.**

| Drug | $I_{max}$ | $T_{50}$ / h | $EC_{50}$ / μM |
|---|---|---|---|
| STS | 2420 | 1.5 | 1.27 |
| Act D | 1280 | 2.0 | 3.66 |

**Supplementary Table 2. Comparison of the anti-cancer efficacyof STS and Act D towards cancer cell and normal cells.**

| Drug | Cell lines | $I_{max}$ | $T_{50}$ / h |
|---|---|---|---|
| STS | HepG-2 | 1970 | 5.2 |
| | L-02 | 2020 | 1.5 |
| Act D | HepG-2 | 1790 | 6.5 |
| | L-02 | 2280 | 2.2 |

**Supplementary Table 3. Comparison of the anti-cancer efficacy of Act D, metformin and their combination evaluated by MTT and scPISA.**

| Drug | MTT | scPISA | | |
|---|---|---|---|---|
| | $IC_{50}$ | $I_{max}$ | $T_{50}$ / h | $EC_{50}$ |
| Act D | 3.16 ± 1.09 μM | 1790 | 6.5 | 1.99 ± 0.29 μM |
| Metformin | 43.50 ±15.09 mM | 600 | 8.8 | 8.34 ± 0.09 mM |
| Act D + Metformin (10 mM) | 2.07 ± 1.38 μM | 2150 | 4.5 | 0.37 ± 0.05 μM |
| Act D + Metformin (20 mM) | 1.61 ± 1.31 μM | 2270 | 4.0 | 0.05 ± 0.002 μM |
